# Fire removes preexisting pyrogenic organic matter from the ecosystem through the mechanisms of both direct combustion and increasing mineralizability

**DOI:** 10.64898/2025.12.04.691661

**Authors:** Mengmeng Luo, Kara Yedinak, Keith Bourne, Thea Whitman

## Abstract

Pyrogenic organic matter (PyOM) produced during fires plays an important role in the global carbon (C) cycle due to its low microbial availability and resulting high persistence. However, PyOM can be combusted or chemically altered in subsequent fires. To explore the effect of subsequent fire on PyOM, we used a mass loss calorimeter to deliver realistic heat fluxes to jack pine (*Pinus banksiana* Lamb.) PyOM produced at 350 °C, testing heat flux profiles (High, Low, and Control) and burial depths (Surface, 1 cm, and 5 cm) in a full-factorial design. We found that both variables significantly affected the C remaining after burns (*p* < 0.05), with greater C losses at higher heat fluxes and shallower exposure depths. Consistent with our predictions, treatments with high heat exposure (HF_High_+Surface, HF_High_+1cm, HF_Low_+Surface) showed increases in pH and decreases in total dissolved organic carbon (DOC) and mineralized C after the subsequent fire. Counter to our predictions, treatments with intermediate heat exposure (HF_High_+5cm and HF_Low_+1cm) caused significant decreases in pH and increases in DOC, mineralized C, modelled decomposable C, and modelled C decomposition rate (*p* < 0.05), compared to the unburned controls. These findings highlight that, despite its high persistence in fire-free conditions, PyOM is readily combusted and altered in subsequent fires, with important implications for changing fire regimes and the global C cycle.

**Synopsis Statement:** Subsequent fires not only consume PyOM produced in previous burns, but can also increase its susceptibility to microbial decomposition, which challenges the potential for PyOM to contribute to long-term carbon storage.

## Introduction

As the second largest pyrogenic carbon (PyC) pool in the world^1,2^, soil PyC is an important component of global carbon (C) stocks. Soil PyC, which can comprise less than 1% to over 45% of total soil organic carbon (SOC)^3^, varies across and within ecosystems, influenced by factors such as land cover, land use, climate, and topography^4^. PyC is a constituent of pyrogenic organic matter (PyOM) – the dark-colored, carbon-rich residue modified during incomplete combustion of carbonaceous materials^5,6,7^. PyC is often characterized by a high degree of aromatic condensation^8,9^, which contributes to its relatively high persistence^10^.

However, PyOM is chemically heterogeneous^1,5,11^ and is certainly not inert. The fate of soil PyOM is influenced by physical, chemical, and biological processes. Its stability is governed by both its intrinsic chemical properties and interactions with the surrounding environment^4,13,14^. Yet, despite growing recognition of its ecological significance, the mechanisms regulating the persistence, transformation, and mobility of PyOM in soil remain insufficiently characterized^15,16,17^.

To characterize the mechanisms that control PyC dynamics, we can turn to the classic soil-forming processes: additions, losses, transformations, and translocations^18^. Each of these processes is significantly influenced by fire, which is naturally a dominant force in PyC cycling^2,8^ and is even proposed to represent a soil-forming factor in itself^19^. Thus, we must also consider each process in the context of “subsequent fire”. Over sufficiently long timescales, almost every fire occurs within the footprint of a previous one. However, patterns of fire activity have been dramatically altered by human influence. In North America, fire exclusion following European settlement disrupted historical fire regimes^20^, while ongoing rising temperatures and increasing drought events have led to greater fire frequencies in many regions^21,22,23^. As a result, fires are now more likely to recur in the same location within ecologically relevant timescales. We refer to these as “subsequent fires” (sometimes termed *reburns* or *repeated fires* in the literature)^7,24^, and within this context, we consider the four processes governing the fate of PyOM in soil.

*Additions:* Fire is the primary and, in many cases, the only natural process responsible for generating new inputs of PyOM in terrestrial ecosystems. Through pyrolysis, fire transforms plant biomass, litter, and soil organic matter (SOM) into PyOM, thereby contributing to the persistent C pool^25,26^. The quantity of PyOM produced is shaped by fire intensity, duration, and fuel type. High-intensity fires, such as crown fires, tend to produce more aromatic and condensed forms of PyC with greater resistance to degradation^5,8^. Spatial and temporal variation in fire regimes also strongly influences PyOM additions across ecosystems. For instance, low-intensity, high-frequency grassland fires may yield smaller PyOM quantities per fire event, but their cumulative contributions over time can be substantial^27^. In contrast, high-intensity, low-frequency forest fires can produce large PyC pulses, though the slow regeneration of woody biomass may limit subsequent additions^28,29^ (Fig. 1a). Thus, PyOM addition rates are closely linked to the frequency, severity, and recovery dynamics of ecosystem fire regimes.

**Figure 1.**
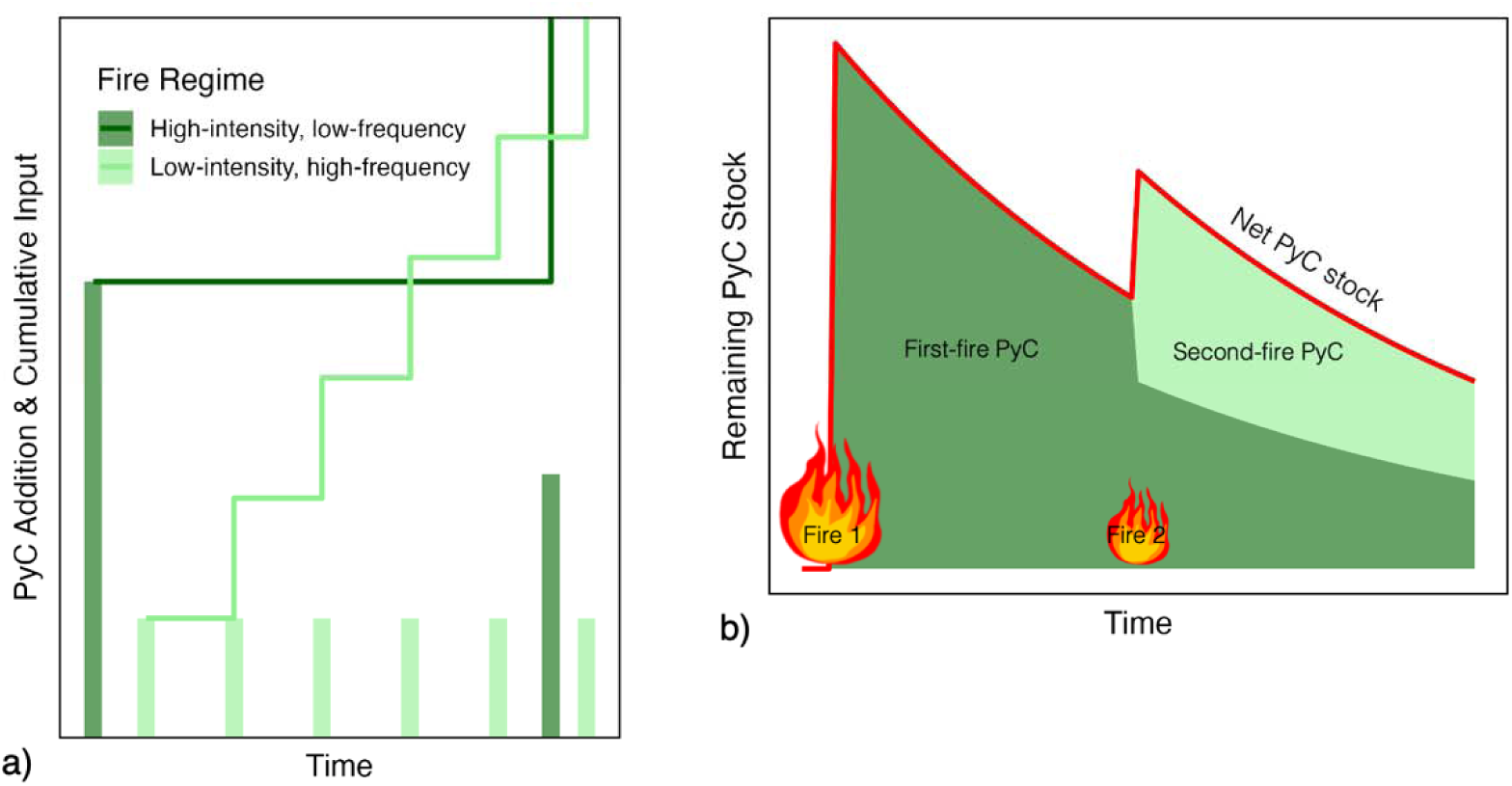
Idealized illustration of fire effects on PyC dynamics. (a) Illustrative PyC additions with each fire event (bars) and cumulative inputs (step lines) under contrasting fire regimes. (b) Conceptual model of PyC stock trajectories with recurrent fires, showing the contributions from first- and second-fire PyC pools (dark and light green, respectively). The red line indicates the net PyC stock over time. (Modeled conceptual trends, not to scale)

*Losses:* PyC can be removed from terrestrial ecosystems through both fast and slow pathways (Fig. 1b). The primary and most immediate loss mechanism is combustion during subsequent fires, in which preexisting PyC is further oxidized to CO_2_. This has been proposed to be the main pathway for the loss of PyC. This fast-acting pathway can result in the complete or near-complete combustion of surface PyC during high-intensity fire events, representing a major disruption to the PyC pool^8,30^.

In contrast, post-fire losses of PyC are considerably slower but long-lasting, with rates decreasing as production temperature increases^12,31^. Chemical and biological oxidation are common mechanisms that lead to PyC loss over time^32,33^. Chemical oxidation, often mediated by oxygen-rich environments or abiotic reactions, can also enhance microbial access to PyC^31^. Microbial decomposition, especially by heterotrophic communities, can slowly mineralize PyC to CO_2_^12,34,35^, though this pathway typically acts on more labile or partially charred fractions. While the rates for slow losses are low compared to the rate of combustion losses, they play an important role in long-term PyC turnover^16,36^.

*Transformation:* The transformation of PyOM during subsequent fires depends not only on reburn conditions, but also on the characteristics of the preexisting PyOM. The degree of condensation and aromaticity can determine whether PyOM is susceptible to further alteration or largely resistant^12^. When exposed to fire again, the outcome is also strongly shaped by oxygen availability and heating intensity. Under aerobic conditions, PyOM may lose more labile or “tarry” fractions, widen pore structures, and increasing surface area and reactivity, thereby enhancing microbial accessibility^37^. In contrast, under anoxic or pyrolytic conditions, condensed structures can be preserved or even further stabilized, increasing resistance to subsequent decomposition^38^.

Beyond fire-driven events, PyOM also undergoes gradual transformation over ecological timescales. Abiotic processes such as physical fragmentation and chemical oxidation, as well as microbial metabolism, can reduce aromaticity and modify surface chemistry^8,39^. Some relatively labile PyC fractions may be assimilated by microbes and subsequently stabilized in microbial biomass or necromass, thereby shifting them into different persistent C pools^32,40,41^.

*Translocations:* During a fire, a small portion of PyC is emitted as aerosol black carbon, which can be transported by wind and later deposited^1,2^. However, the bulk of PyC typically remains on the surface or in the soil, where it can be translocated through both lateral (e.g., surface run-off, aeolian transport) and vertical (e.g., leaching, bioturbation) pathways. These post-fire PyC movements remain poorly quantified^1,2,4^.

As with total SOC, PyC concentration typically decreases with soil depth and has a strong positive correlation with total SOC concentration^42^. Evidence also suggests that PyC may be preferentially transported downward relative to other forms of SOC^17^. Additionally, small particle sizes (<53µm) or bigger surface areas facilitate the adsorption of PyC onto fine soil particles and enhance downward movement^43,44^. Finer, older, and more aromatic PyC is often found in deeper soil profiles, suggesting that water percolation, interacting with other transformation processes, contributes to vertical redistribution over time^17,45^. However, this process is slow in nature and may be inhibited by fire-induced hydrophobicity, which limits water infiltration^46^, at least in the short term post-fire.

Lateral transport also plays an important role, particularly through post-fire erosion. Water and wind can relocate PyOM from its point of origin and deposit it in topographic depressions^47^, water bodies, or seasonally flooded anaerobic sites, thereby reducing microbial decomposition and enhancing persistence^15^. In rare cases, such depositional events, especially burial by loess, can preserve PyC for millennia in deep, oxygen-limited environments^48^.

Despite these potential pathways, the majority of PyC tends to remain near the soil surface. Over decadal timescales, more than 70% is found in the top 10 cm, and most within the top 30 cm^49,50,51^. In shorter-term studies following recent PyOM deposition, PyC is typically concentrated in the top 15 cm^35,52^. These consistent surface accumulations highlight the relatively low mobility of PyC under most field conditions, despite the range of processes that can facilitate its translocation.

Together, additions, losses, transformations, and translocations control the size and nature of PyC stocks in soils, and are, in turn, influenced by subsequent fire. Changes in the Anthropocene further increase the need to understand how PyC produced during one fire is affected by a subsequent fire in the context of these processes. In a previous companion study, we designed a system to deliver realistic heat fluxes to a pyrocosm to simulate subsequent fire effects on PyOM in a sand matrix^53^. Building on this system, our present study addresses the central question: How does subsequent fire influence the chemical properties (transformation) and the microbial decomposition (losses) of the preexisting PyC (Figure 2)?

**Figure 2.**
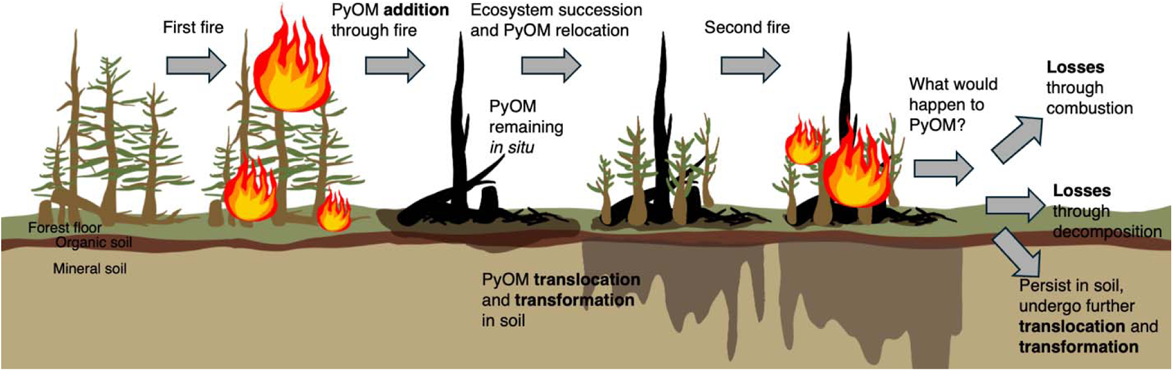
Conceptual diagram illustrating our central research question: What will happen to PyOM during and after a second fire? After the first fire, the residual PyOM on the forest floor and in organic soil decreases due to translocation and microbial degradation. Some PyOM percolates into mineral soil with water and biological activity and is deposited deeper in the soil profile^1,2,35^. This diagram drew inspiration from Fig. 2 in Santín et al. (2016)^2^.

This study focuses on three main aspects of PyC persistence during and after reburns: 1) protection through burial at different depths (effects of previous translocations); 2) changes in potential for leaching and microbial availability, assessed via dissolved organic carbon (DOC) (transformations); 3) changes in biological degradation, assessed through cumulative C mineralization (losses). We hypothesized that the combination of different heat fluxes (HFs) and depths would influence pH, total C content, DOC, and the mineralized C of the PyOM in the subsequent fire. Higher HFs and shallower burial depths were expected to result in higher pH, due to greater ash deposition on and near the surface. However, these same conditions would lead to lower total C, mineralized C, and DOC, reflecting increased thermal degradation and reduced microbial accessibility.

## Materials and Methods

We designed our system to represent a jack pine-dominated pine barrens (*Pinus banksiana* Lamb.) characteristic of Wisconsin, USA. Pine barrens typically have well-drained to excessively well-drained sandy soils, and the natural fire return interval for jack pine stands in the area is 3-7 years^54^, generally with low fire severity. We produced PyOM from jack pine using a modified muffle furnace^53,55^ at 350 °C (77.76 ± 0.12% of C), using wood from a small jack pine tree from Wilson State Forest Nursery (43.1461124, -90.6950388).

We simulated the effects of fire using the method described in detail in Luo et al. (2025)^53^. Briefly, we used a mass loss calorimeter to expose PyOM to high and low heat fluxes (HF_High_ and HF_Low_) at the surface, buried 1 cm, and buried 5 cm deep in an ashed sand matrix (“pyrocosm”), chosen to represent the typically sandy soil of jack pine forests. The high and low heat fluxes, representing different heat received on the soil surface, were parameterized using data from log burns^53^. For each depth and heat treatment, we had five individual replicates (1 g PyOM mixed with 8 g sand), which were exposed to the same heat flux (or control) at the same time, but were distanced from each other during exposure and then analyzed individually (Fig. 3). We measured chemical parameters including pH, total C and N, and DOC, and evaluated biological C mineralization, to test how different heat exposures affect the chemical and biological properties of PyOM during a simulated subsequent fire.

**Figure 3:**
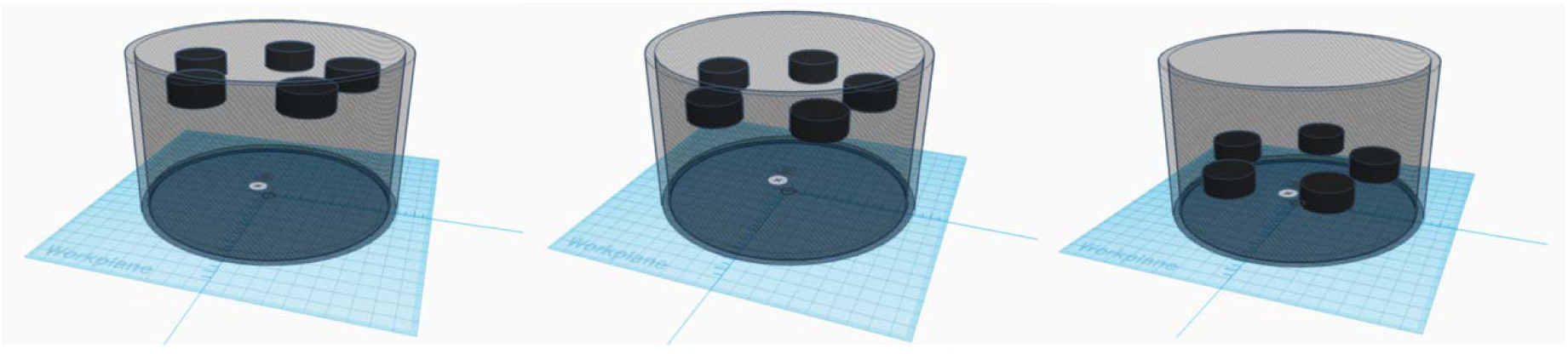
Relative locations for the PyOM samples (from left to right: Surface, 1 cm, 5 cm; figures generated from Tinkercad)

### Sample collection, unified dilution, and homogenization

In order to ensure we captured as much of the PyOM as possible, we collected and analyzed it as a PyOM-sand mixture, rather than trying to physically separate the PyOM from the sand matrix. By analyzing blank sand along with sand-PyOM mixtures, we can infer the properties of PyOM without overlooking tiny particles that are difficult to retrieve, or soluble components that would be lost if we had used a water or other density-based extraction. For the total elemental analyses, this should have no effect on our calculation of stocks. For the pH measurements, we would expect different absolute values, but observe the same trends with treatment that we would see if it were possible to separate and analyze PyOM without sand. For the C mineralization assays, we would again expect different absolute values, but the same trends.

All PyOM-sand matrix samples were collected from the pyrocosm after undergoing sufficient cooling to be safe to handle. For 1 cm and 5 cm samples, before retrieving the samples, the sand on the top was carefully scraped away down to just above the depth of the samples. After the PyOM retrieval, we used a custom-designed 3D-printed PLA sample divider (Fig. 4) (modeled with Tinkercad; printed with Ultimaker S5, UW Makerspace) to collect the rest of the sand (along with any trace amounts of remaining PyOM) in the pyrocosm. The sample divider includes a sharp cutting end with five wedges (with interior angle of 72°) that allows it to evenly separate the samples by cutting through the sand to the bottom of the pyrocosm. Then we flipped the divider and the pyrocosm together. The collection area at the other end has five identical chambers to hold the sand in individual sections for each replicate (Fig. 4), so the sand drops into each chamber. To retrieve the sand from each chamber, we poured the sand out from one chamber while covering the other chambers at the same time. For the unburned controls, PyOM was buried as for the burn treatments, and then retrieved using the same processes as for the burned samples.

**Figure 4.**
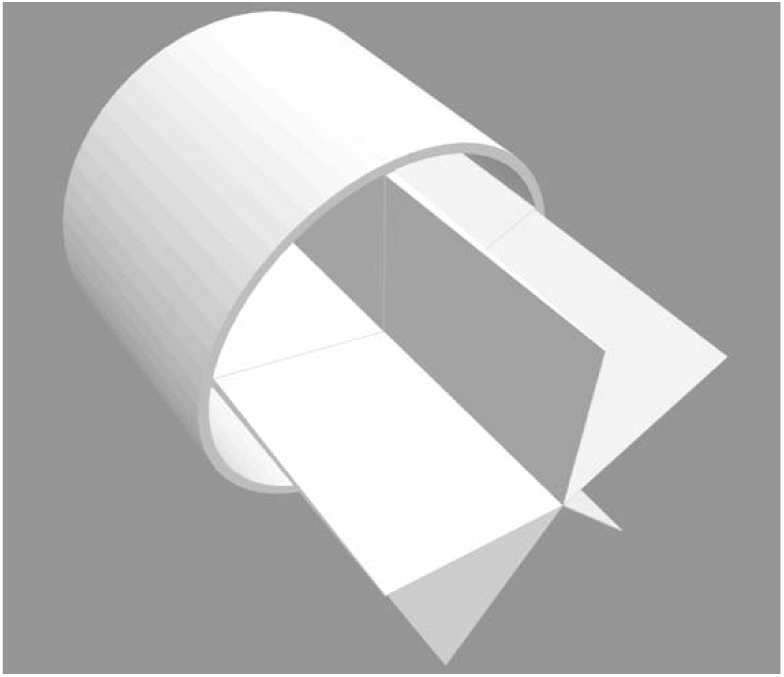
Sample divider. The divider has five identical wedges with 72° interior angles housed in a cylinder container with an interior diameter the same as that of the pyrocosm, creating five identical chambers in the cylinder container to collect sand; modeled in Tinkercad.

After initial collection, the mass of each burned and retrieved PyOM-sand sample was recorded. Before homogenizing the samples, we ensured there was the same mass of sand in each PyOM-sand mixture. To do this, we calculated the mean mass loss for all samples in the same treatment (assuming the mass loss for the whole pyrocosm is, then the mean mass loss is ,)then subtracted it from the original PyOM mass to get the mean remaining PyOM mass in each sample, allowing us to calculate the mass of sand in each mixture. Then we added the right amount of sand (using reserved sand from the corresponding section of the pyrocosm where each sample was located) to the sand-PyOM mixture required to bring the final mass of the sand to 80 g for each sample. Thus, all final samples had 80 g sand mixed with whatever mass of PyOM out of the initial 1 g remained. For further analysis, all the samples were ground and homogenized individually in Fritsch Vibrating Cup Mill Pulverisette 9 at 1500 rpm for 10 seconds. We also homogenized a subsample of ashed and washed sand, which was used as the sample blank for subsequent analyses. All of the following analyses were done using these ground and homogenized samples.

### Measuring pH

For each sample (PyOM-sand mixture) and the sample blank (ground quartz sand), 0.5 g was weighed in a 15 mL Falcon tube and saturated with 0.5 mL of DI water for 48 hours before the measurement. Because the samples were dry, the initial addition of water likely initiated various chemical reactions within the sample that led to unstable pH values immediately after wetting-up. To allow the sample to equilibrate at a stable pH, we let them sit for 48 hours before measuring pH. Ecologically speaking, this wet-up process could roughly represent the first rain after a fire. After the equilibration, another 9.5 mL of water was added to each tube, bringing the solid (g) to liquid (mL) ratio to 1:20, following Zeba et al. (2022)^56^. Then all tubes were oscillated in an INCU-SHAKER 10L (Benchmark Scientific, Sayreville, NJ, USA) for 10 minutes at 200 rpm and centrifuged with a Centrifuge 5810 R (15 amp version, Eppendorf, Hamburg, Germany) at 3214 rcf for 15 min. Two lab replicates of 1 mL of the supernatant solution were extracted from the tube for each sample and measured with a pH electrode InLab Micro (Mettler Toledo, Columbus, OH, USA) connected to a Thermo Scientific Orion Star A215 pH/Conductivity Meter (Thermo Fisher Scientific, Waltham, MA, USA). Lab replicates were subsequently averaged.

### Total carbon

Samples used for measuring total C were dried at 60 °C for 48 hours before preparation. Samples were wrapped in CE Elantech 5 x 9 mm tin capsules (∼60 mg, for treatments HF_High_+5cm, HF_Low_+1cm, HF_Low_+5cm, and controls) and 10 x 12 mm tin capsules (200 ∼ 400 mg, for treatments HF_High_+Surface, HF_High_+1cm, and HF_Low_+Surface, which had lower C concentrations) and analyzed using a Thermo Flash EA 1112 NC Analyzer (Thermo Fisher Scientific, Waltham, MA, USA) at the Jackson Lab, UW-Madison. Two lab replicates were analyzed for each sample and averaged. Total C is presented as C stocks (mg of C per sample).

### Dissolved organic and inorganic carbon

DOC samples were prepared with a 1:3 sample-to-water ratio using DI water (7 g sample with 21 mL DI water). Samples were oscillated at 150 rpm for 60 minutes and centrifuged at 2000 rcf for 10 minutes^57,58^. The supernatant of the solution was extracted with a syringe and filtered through the GD/X 25 mm Sterile Syringe Filter (glass microfiber filtration medium, 0.45 µm) into 17 mL TOC tubes (ashed at 450 °C for 8 hours in a muffle furnace to remove any trace C). All samples were analyzed within 24 hours of the extraction and stored at 4 °C before analysis. Total dissolved carbon (TC) and dissolved inorganic carbon (DIC) were measured by combustion using a Sievers M5310 C Laboratory TOC Analyzer System with GE Autosampler (General Electric, Boston, MA, USA) at the Water Science and Engineering Laboratory, UW-Madison. DOC was calculated by subtracting DIC from TC. The raw DOC data were translated into units of “mg DOC per PyOM-sand sample” using Equation 1,

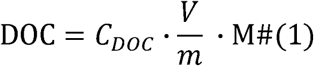

where *C_DOC_* is the concentration of DOC in mg L^-1^, which is the direct measurement from the equipment. *m* is the mass of sample (7 g) that is used for DOC extraction. *V* is the volume of DI water added to sample (0.021 L). M is the total mass (∼80 g) of each sample. DIC was also calculated with the same equation.

### Twelve-week incubation

To investigate PyC mineralization after subsequent fire, we incubated samples for 12 weeks with a microbial inoculum. The setup for the 12-week incubation was comprised of a vial of 15 g sample with microbial inoculant added and a vial of KOH solution (the CO_2_ trap) placed inside a sealed 8 oz (236.6 mL) glass jar (Burch Bottle & Packaging)^59^. All samples, along with 3 sample blanks (ground quartz sand) were included in the incubation.

We inoculated each sample with a microbially active soil extraction along with a nutrient solution, with the aim of avoiding nutrient limitation and focusing on C mineralizability. The inoculant for the sample incubation was composed of a microbial extraction from a previously burned soil, collected in 2020, six years after the 2014 King Fire (38.86953, -120.61322) in California^60^. The soil is a sandy clay loam, with a pH of 5.7. After the soil was retrieved from the -23 °C freezer and thawed at 4 °C, 5 g of the soil was incubated at 25 °C in a VWR Incubator Gr Con 4CF (Thermo Electron, Langenselbold, Germany) at 60% water holding capacity (WHC) for 8 days in order to reactivate the microbial community^61^. This resulting community would of course not be expected to perfectly mimic a natural soil microbial community – rather, our goal was to derive a complex community that would be likely to have natural abilities to degrade PyOM, given its recent fire history.

To create the inoculum, 4.5 g incubated soil was added to 225 mL of DI water (1:50 w/v) and shaken at 100 rpm for 30 minutes. Then the solution was left to settle for 10 minutes and filtered through Whatman 1 filter paper. This extract was mixed with a nutrient solution in a 500 mL volumetric flask (with DI water added to 500 mL). The resulting 500 mL nutrient-inoculant solution contained the microbial extract, 4 mM NH_4_NO_3_, 4 mM CaCl_2_, 2 mM KH_2_PO_4_, 1 mM K_2_SO_4_, 1 mM MgSO_4_, 25 µM H_3_BO_3_, 2 µM MnSO_4_, 2 µM ZnSO_4_, 2 µM FeCl_2_, 0.5 µM CuSO_4_, and 0.5 µM Na_2_MoO_4_^61,62^.

Each 15 g PyOM-sand sample received 2.05 mL of nutrient-inoculant solution, representing 55% WHC, and was placed in the larger glass jar. 5 mL of CO_2_-depleted DI water was added to the bottom of each jar to maintain a moist environment during the incubation^63^. Each jar received a vial with 15 mL 0.05 M KOH to serve as the CO_2_ trap, and was sealed for incubation at 25 °C in a VWR Incubator Gr Con 4CF (Thermo Electron, Langenselbold, Germany).

For each sample, the vial of KOH solution was retrieved and replaced with a new vial of KOH solution 2 weeks, 4 weeks, 8 weeks, and 12 weeks after the start of the incubation. At each retrieval, the electric conductivity (EC) of the KOH vial was measured with an Orion DuraProbe Conductivity Probe connected to the Orion Star A215 pH/Conductivity Meter (Thermo Fisher Scientific, Waltham, MA, USA). The EC of the new KOH solution was also recorded at the beginning of the incubation and during each retrieval.

The KOH trap is used because KOH reacts with CO_2_ to produce K_2_CO_3_ (Equation 2), which causes a linear change in EC. Based on the change of EC, we can calculate how much CO_2_ is emitted^63^ (Equation 3).

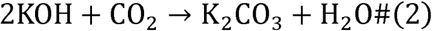

We also ran a “blank incubation” to evaluate how much CO_2_ is emitted from the dissolution of inorganic C (especially for the treatments with higher ash production). This set of incubation trials was inoculated with only nutrient solution (substituting the microbial extract with DI water) and filtered through a 0.2 µm cellulose acetate sterile syringe filter to minimize the potential for contamination by microbes.

### Carbon mineralization

The calculation of C loss as CO_2_ (CO_2_-C, in mg) in each PyOM-sand sample follows Equation

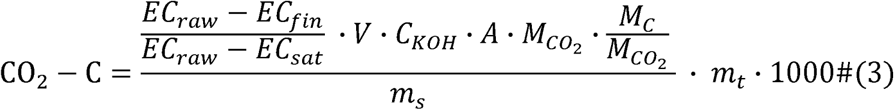

where *EC_raw_* is the EC measured for pure 0.05 M KOH. *EC_sat_* is the EC measured for 0.025 M K_2_CO_3_ (the result when all OH ions have reacted with CO_2_). *EC_fin_* is the EC measured at each retrieval. *V* is the volume of the KOH trap in liters (0.015 L). *C_KOH_* is the molarity of the KOH trap (0.05 M). *A* is the ratio of CO_2_ needed to react with one portion of KOH (0.5). is the molar mass of CO_2_ (44.01 g mol^-1^). is the molar mass of C (12.011 g mol^-1^). is the mass of sample used for incubation (∼15 g). is the total sample mass for each sample (∼80 g). Measurements from the sample blanks (quartz sand with inoculant) were averaged and deducted from each sample to account for CO_2_ present in the jars when they were sealed and any trace amounts of C from the quartz sand and inoculant itself.

In order to compare the relative mineralizability of the samples, we also reported the fraction of total post-heating C (mg CO_2_-C per mg total C) respired as CO_2_ during the entire 12-week incubation (C_min_, Equation 4), where C_%_ is the percentage C for each sample and is the total sample mass for each sample (∼80 g).

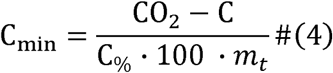

Fraction of C remaining (C_r_) after mineralization was calculated with Equation 5.

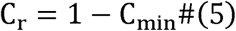

### Modelling mineralizable carbon and carbon decomposition rate

A one-pool decay model (modified from Singh et al., 2012^16^) (Equation 6) was used to fit the values of C_r_ (Equation 5) for each treatment using nlsLM function in minpack.lm package in R^65^,

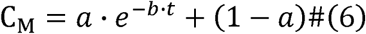

where *C_M_* is the modeled fraction of C remaining, coefficient is the readily decomposable fraction of C, and coefficient is the decomposition rate (per week), and is the time (in weeks) after the start of incubation. The first part of the equation represents a typical decay model. Because PyC is so persistent, a simple exponential decay model did not fit the data well, so we added the second term (1-a) to represent the fraction of C that is relatively non-decomposable (over the duration of the incubation).

### Statistical analyses

Calculations were performed in Excel and R^66^. A segmented regression model^67^ was used to fit the peak temperature as the predictor for pH. The purpose of this model is to determine what temperature is associated with the lowest pH values. A two-way ANOVA with an interaction term for heat flux and burial depth, with Tukey’s HSD^68,69^ was used to determine if there were significant differences between treatments for pH, total C, mineralized C, DOC, and DIC. A oneway ANOVA and Tukey’s HSD^68,69^ were used to determine if there were significant differences between treatments for C mineralizability, decomposable C fraction (coefficient ), and C decomposition rate (coefficient ). Linear models^70^ were used to assess correlations between 1) decomposable C fraction vs. C mineralizability, 2) DOC and DIC vs. pH, and 3) DOC vs. mineralized C. For some treatments (HF_High_+Surface and HF_High_+1cm), total remaining C was below detection limits, so these treatments were excluded from analyses of C mineralization and mineralizability. Potential outliers from mineralized C, C mineralizability, and decay model were determined by *z*-score or iterative *z*-score and removed from analyses if |*z*| > 3 (3 standard deviations away from the mean). All the figures for visualizing data were made using ggplot2 in R^71^.

## Results

### As heat exposure increased, pH first decreased, then increased

Both HF and burial depth significantly affected pH (*p* < 0.001, ANOVA in Table S1.2, Tukey’s HSD, Fig. 5a), with pH initially decreasing as peak temperature increased, but then increasing as peak temperature reached its highest values (Fig. 5b). Compared to the control, the pH was significantly higher at the surface and at 1 cm under the high heat flux treatment, and significantly lower than the control for the treatments that resulted in intermediate heat exposure (HF_High_+5cm and HF_Low_+1cm). PyOM was not significantly different from the controls for the low heat flux at the surface and at 5 cm.

**Figure 5.**
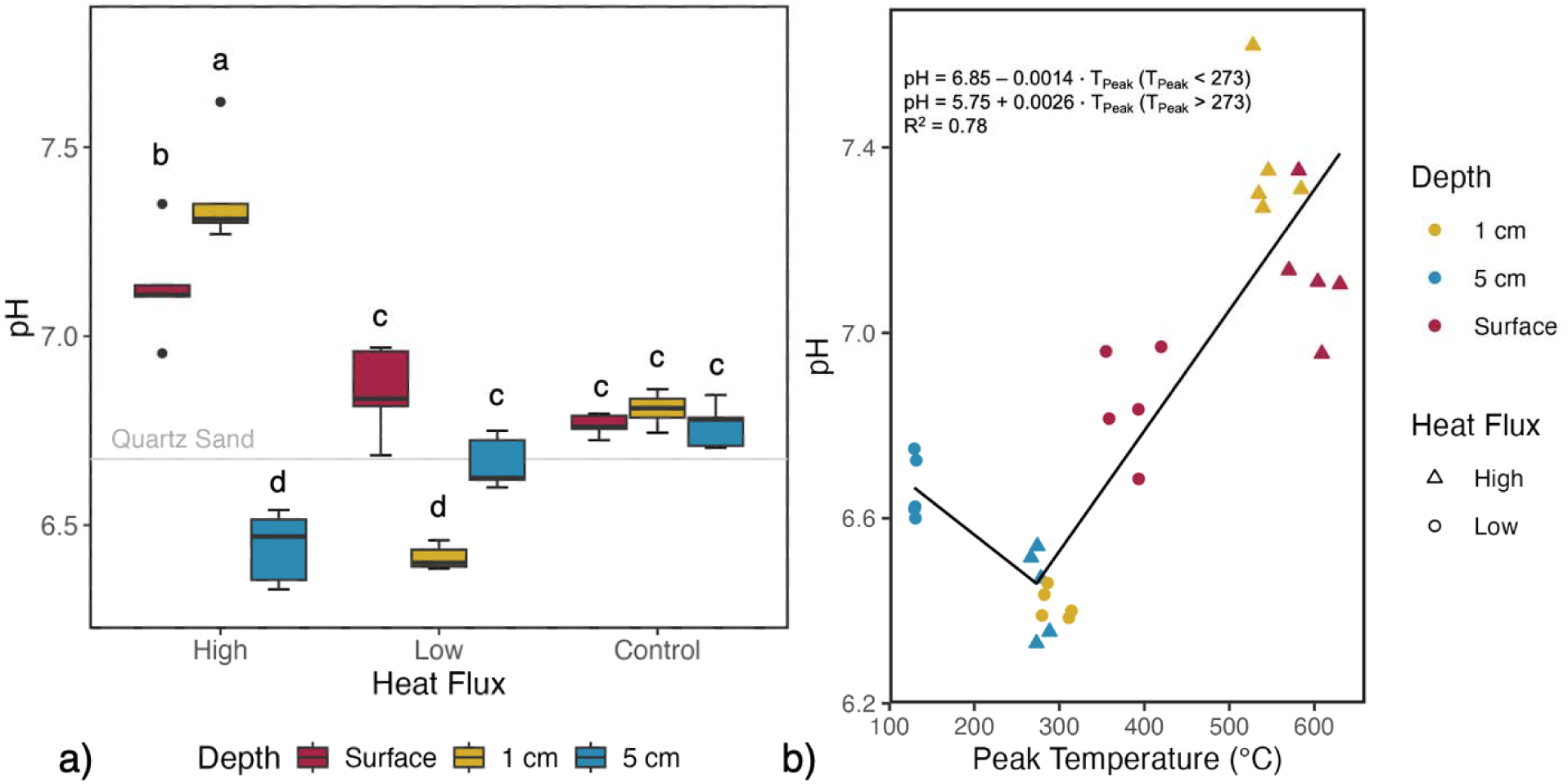
a) pH of each treatment (n = 5, *p* < 0.001; grey line indicates pH value of sample blank; letters indicate significant differences; ANOVA, Tukey’s HSD). b) pH vs. the peak temperature of PyOM (lines and equations indicate fits for segmented regression model; R^2^ = 0.78)

The segmented regression model fit the pH vs. peak temperature data relatively well (R^2^ = 0.78), and indicated that peak temperatures of 273 °C were associated with the lowest pH values (Fig. 5b). Before the breakpoint (273 °C), peak temperature did not significantly affect pH, and after the breakpoint, increasing peak temperatures were associated with increases in pH (Table S1.1 and Text S1.1).

### Carbon loss increased with heat flux and decreased with depth, with near-complete combustion at the highest heat exposure

At the same exposure depth, HF_High_ caused more C loss than HF_Low_, while under the same HF, C loss decreased as burial depth increased (Fig. 6). The C concentrations of HF_High_+Surface and HF_High_+1cm treatments were below the quantification limit (< 0.05%, Fig. 6), indicating near-complete combustion. Almost all the samples also had N contents below the quantification limit (< 0.03%) except for a few controls, so the data for N concentrations are not presented. In general, both HF and burial depth significantly influenced the C loss of PyOM (*p* < 0.001, ANOVA in Table S2.1, Tukey’s HSD, Fig. 6).

**Figure 6.**
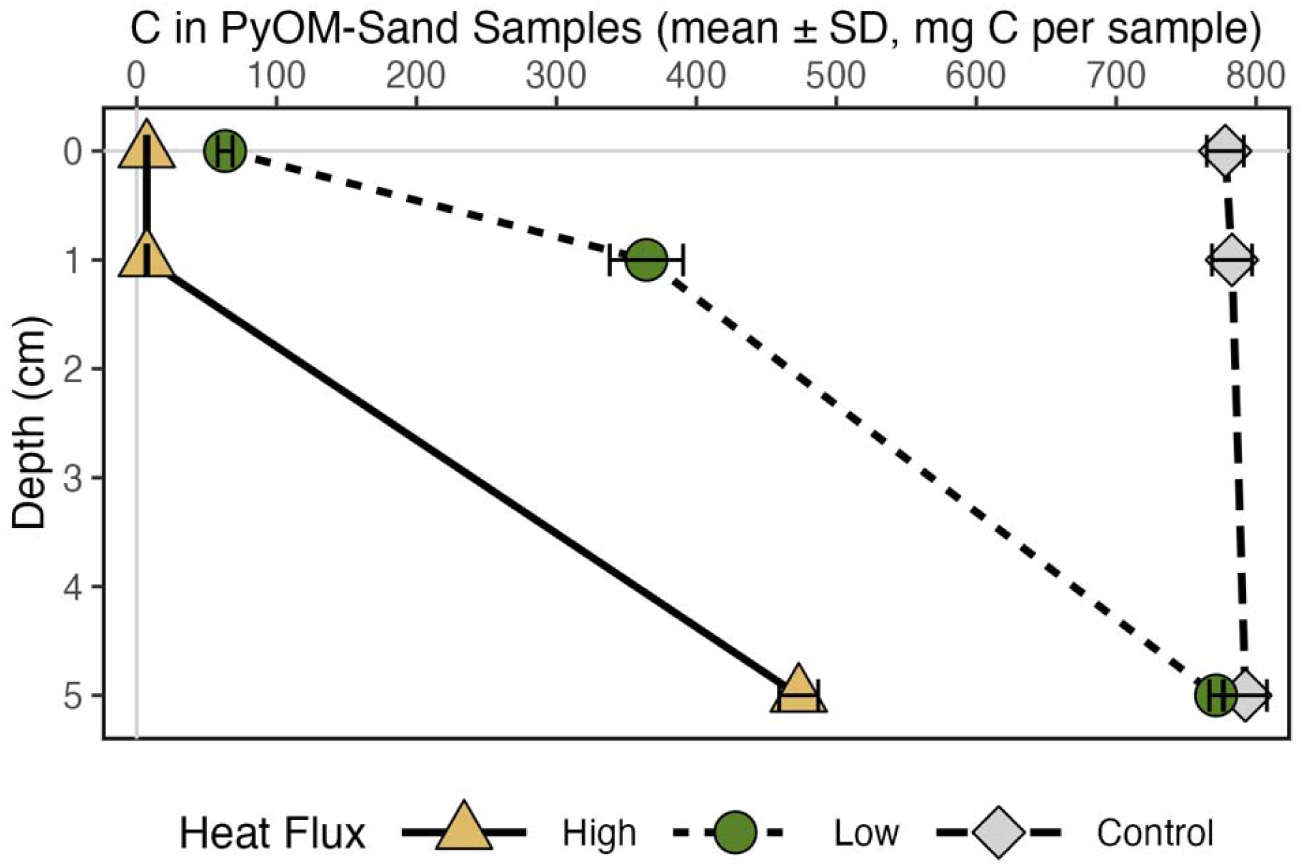
Mass of C remaining (n = 5, error bars represent ± standard deviation. Depth and treatment are both significant, *p* < 0.001; ANOVA).

### Intermediate-to-high heat exposure increased PyOM susceptibility to loss via microbial decomposition after fire

Counter to our predictions, intermediate heat exposure (HF_High_+5cm and HF_Low_+1cm) generally resulted in the greatest total C mineralization during the 12-week incubation (Fig. 7a) and the highest relative C mineralization rates (on a per g of post-heating C basis; Fig. 7b). These treatments had greater C mineralization than those with low or no heat exposure (HF_Low_+5cm and the controls). Highest heat exposure (HF_High_+Surface, HF_High_+1cm, and HF_Low_+Surface) resulted in less C mineralization than HF_Low_+5cm and controls (*p* < 0.05, ANOVA in Table S3.1, Tukey’s HSD, Fig. 7a). One sample from the HF_High_+Surface treatment had extremely high respiration measurements (*z* = +5.42), possibly due to a leak in the jar or sample contamination, and was thus excluded from statistical analyses and figures throughout.

**Figure 7.**
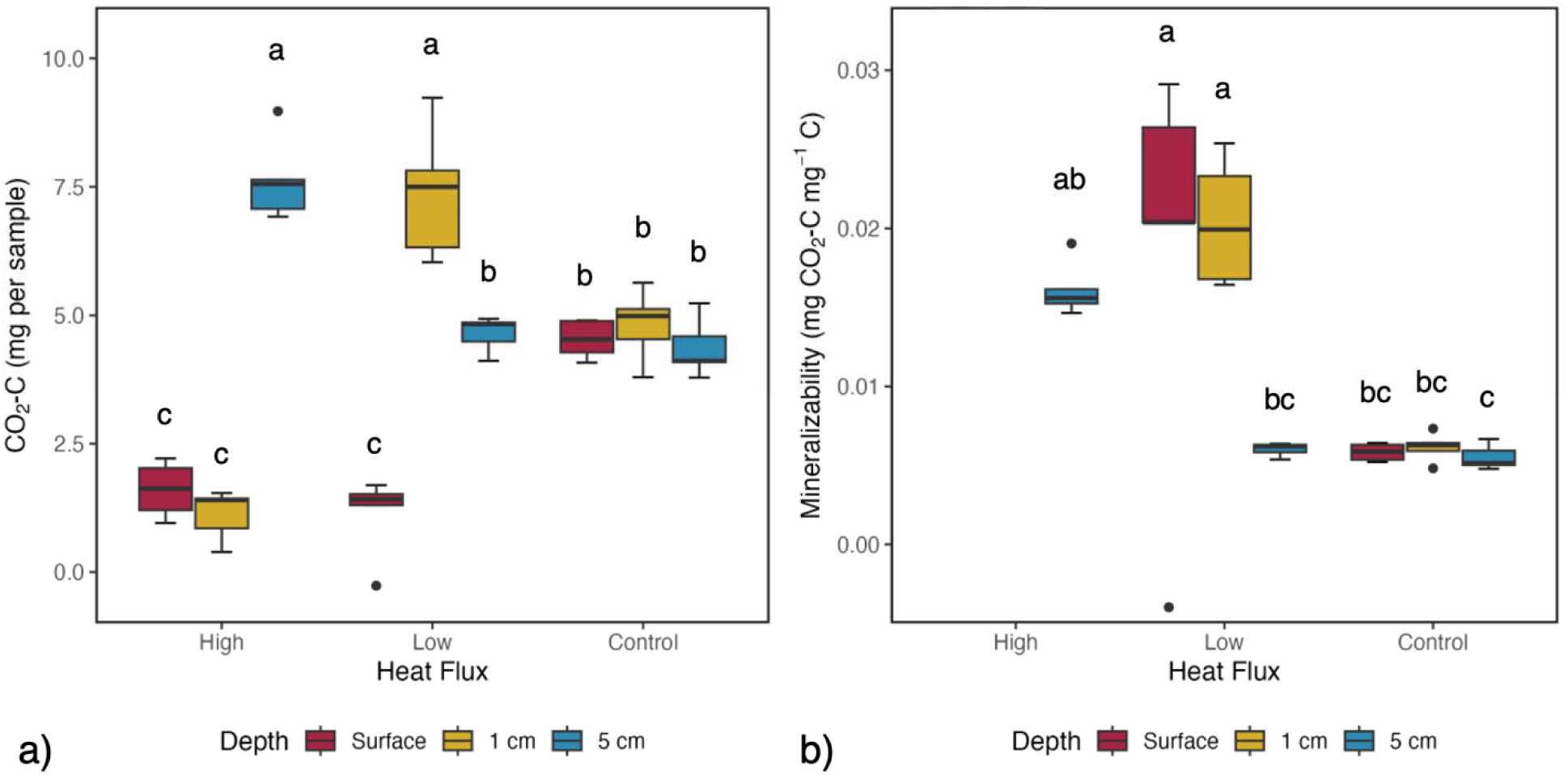
a) C loss as CO_2_ during 12-week incubation (n = 5; one outlier (*z* = 5.42) removed from HF_High_+Surface; letters indicate significant differences across all treatments (*p* < 0.05; ANOVA, Tukey’s HSD)); b) Mineralizability of PyC (n = 5; Treatments HF_High_+Surface and HF_High_+1cm were excluded due to undetectable amounts of total C; letters indicate significant differences across all treatments (*p* < 0.05; ANOVA, Tukey’s HSD))

Mineralizability was higher in treatments with intermediate heat exposure (HF_High_+5cm, HF_Low_+Surface, and HF_Low_+1cm) than HF_Low_+5cm and the controls (*p* < 0.001, ANOVA in Table S4.1, Tukey’s HSD, Fig. 7b). Higher heat exposure might also lead to increased C mineralizability, but C mineralization per g remaining C could not be calculated because total C was below detection limits.

Over time, CO_2_ production generally declined, with most respiration occurring in the first few weeks. Fits for the one-pool decay model of remaining C during the 12-week incubation were good for most of the samples (Fig. S5.1, Table S5.4, R^2^ typically > 0.95). We excluded one sample in the HF_Low_+Surface treatment (Fig. S5.1, top middle panel) because both coefficients in the model were outliers (iterative *z* = -3.93 for coefficient *a*; iterative *z* = 10.30 for coefficient *b*), compared to the rest of the samples in the same treatment. This was due to a presumably erroneous negative C mineralization value measured on Week 12.

The readily decomposable C fraction was highest for HF_Low_+Surface, followed by the intermediate heat exposure treatments (HF_High_+5cm and HF_Low_+1cm), and then HF_Low_+5cm and the controls (*p* < 0.001, ANOVA in Table S5.2, Tukey’s HSD, Fig. 8a). This modelled parameter was significantly and positively correlated with sample mineralizability (Fig. 8c, Table S5.1). Decomposition rate was higher than controls for HF_Low_+1cm, with small or non-significant differences in all other treatments (*p* < 0.001, ANOVA in Table S5.3, Tukey’s HSD, Fig. 8b).

**Figure 8.**
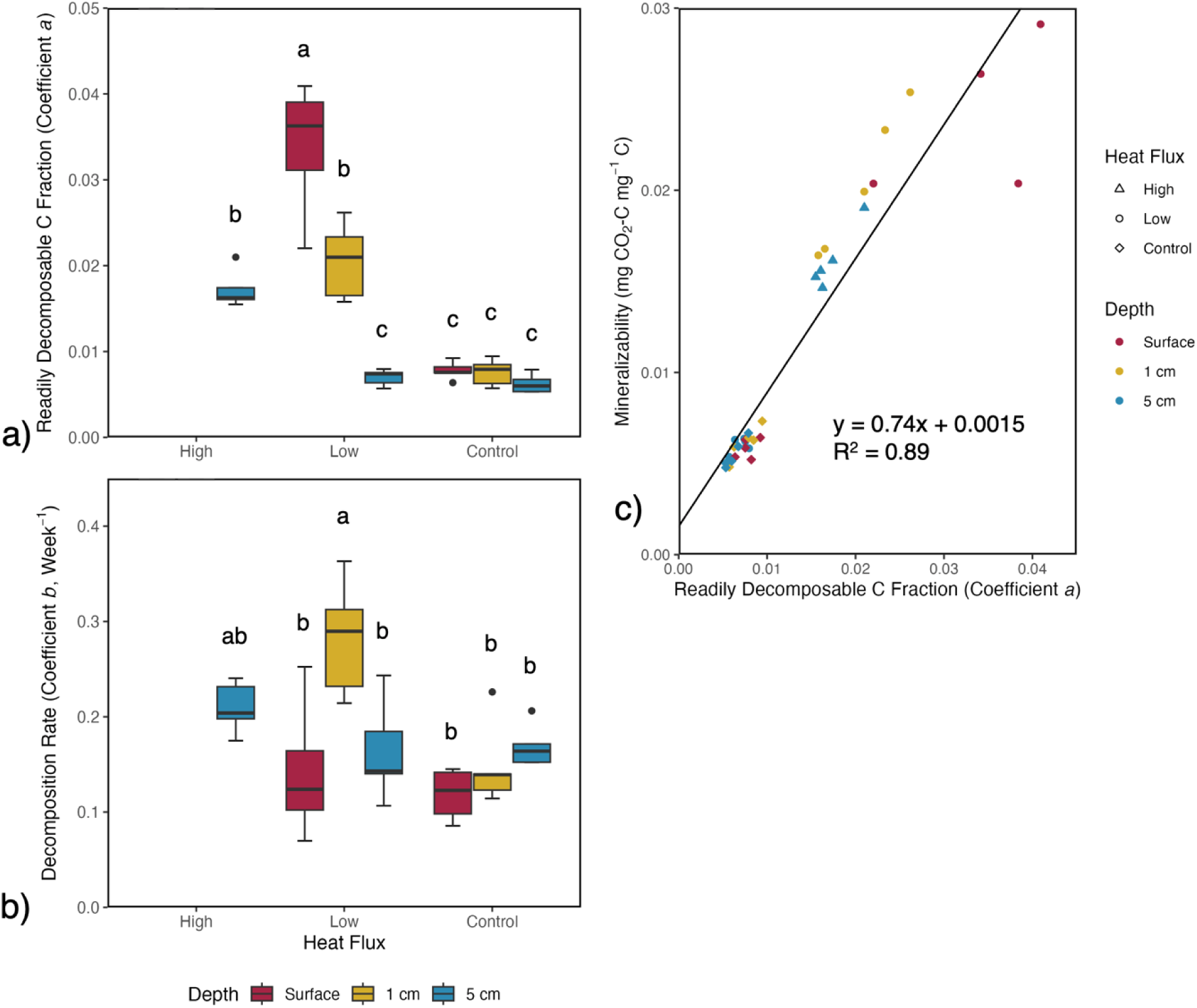
a) Readily decomposable C fraction (coefficient *a*) of the samples from the decay model (n = 5; *p* < 0.001; excluded one outlier in Low HF + Surface); b) Decomposition rate (coefficient *b*) of the samples from the decay model (n = 5; *p* < 0.001; excluded one outlier in Low HF + Surface); c) Mineralizability vs. Readily decomposable C fraction (coefficient *a*) (line indicates linear fit; *p* < 0.001 for slope; R^2^ = 0.89)

### More DOC is produced from preexisting PyOM after intermediate heat exposure

DOC significantly increased in treatments with intermediate heat exposure (HF_High_+5cm and HF_Low_+1cm) compared to the controls (*p* < 0.001, ANOVA in Table S6.4, Tukey’s HSD, Fig. 9a). For the treatment HF_Low_+1cm, DIC significantly decreased, compared to the controls (*p* < 0.001, ANOVA in Table S6.5, Tukey’s HSD, Fig. 9b). In treatments with higher heat exposure

**Figure 9.**
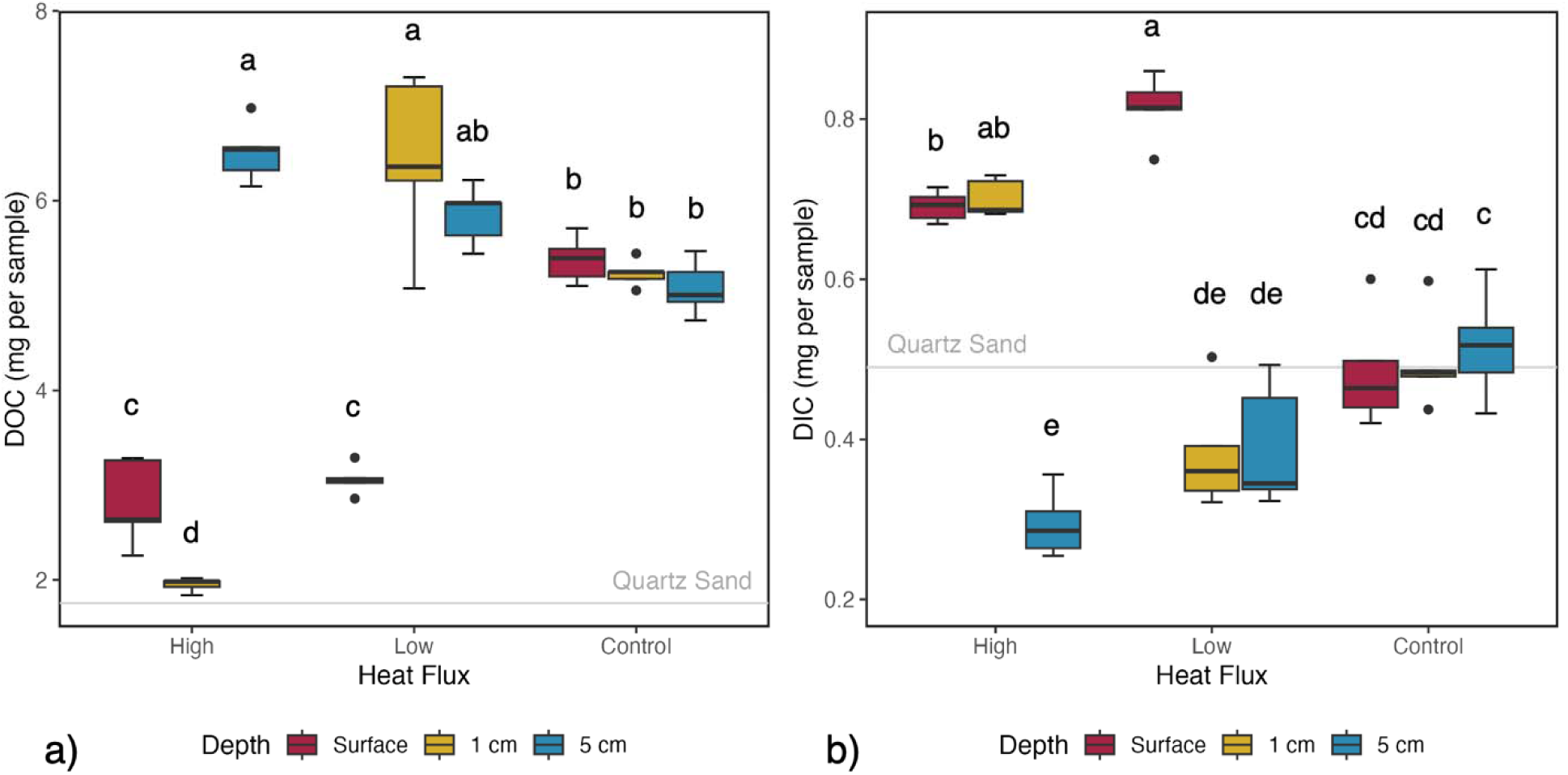
a) DOC of samples. Letters indicate significant differences (ANOVA, Tukey’s HSD, *p* < 0.001); b) DIC of samples. Letters indicate significant differences (ANOVA, Tukey’s HSD, *p* < 0.01). Grey line indicates sample blank. Mean value for sample blanks with only ground quartz sand is indicated with a grey line.

(HF_High_+Surface, HF_High_+1cm, and HF_Low_+Surface), DOC significantly decreased (*p* < 0.001, ANOVA in Table S6.4, Tukey’s HSD, Fig. 9a) while DIC significantly increased (*p* < 0.001, ANOVA in Table S6.5, Tukey’s HSD, Fig. 9b).

DOC was positively correlated with total mineralized C after 12 weeks’ incubation (R^2^=0.83, Fig. 10a, Table S6.1). In addition, pH was negatively correlated with DOC concentrations (R^2^=0.78, Fig. 10b, Table S6.2) while it was positively correlated with DIC concentrations (R^2^=0.54, Fig. 10c, Table S6.3).

**Figure 10.**
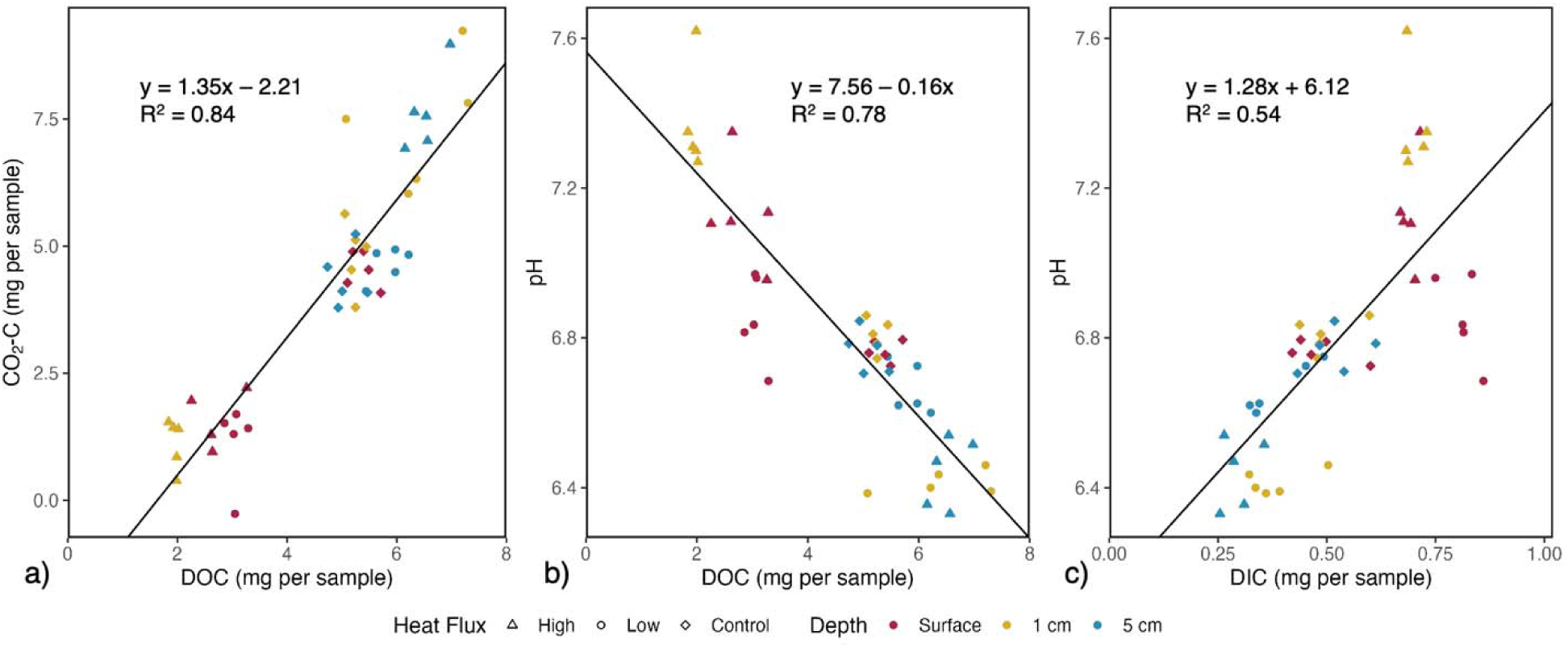
a) CO_2_-C vs. DOC (line indicates linear fit; *p* < 0.001 for both slope and intercept ; R = 0.83); b) pH vs. DOC (line indicates linear fit; *p* < 0.001 for both slope and intercept; R^2^ = 0.78); c) pH vs. DIC (line indicates linear fit; *p* < 0.001 for both slope and intercept; R^2^ = 0.54).

## Discussion

### Heating generally increased pH, but intermediate heating decreased pH

In this study, subsequent fire both decreased and increased the pH of PyOM, depending on heat flux and depth. Contrary to our initial hypothesis of a consistent increase, we observed a nonlinear pattern. While counter to common assumption that pH inexorably increases with fire, these results are similar to past soil-heating studies (samples that are not PyOM-only): pH decreased after low-temperature heating (150-250 °C) before rising again at higher temperatures^72,73^. Our segmented regression model indicated that a peak temperature of 273 °C was associated with the lowest pH (Fig. 5b), which was a little higher than the range reported for soils, suggesting that intermediate heating can produce transient acidification.

The pH decrease at lower temperatures may be linked to volatilization and condensation of organic acids during heating and subsequent cooling, consistent with our observation that higher DOC coincided with lower pH (Fig. 10b). In contrast, pH increases at higher temperatures can result from the release of alkaline cations (K^+^, Ca^2+^, and Mg^2+^) during combustion, and from the loss of proton-donating functional groups^72,74,75^. This interpretation aligns with the positive correlation we observed between pH and DIC, which should reflect carbonate/bicarbonate ions and associated base cations^76^.

At the lowest heating levels (<150 °C), we found little change in pH, in agreement with Fernández et al. (1997)^73^. However, measurable mass losses^53^ and C losses indicate that chemical transformations likely still occurred, although not enough to alter pH.

In general, our results reinforce past studies showing that pH does not invariably rise with fire or heating. Instead, intermediate heating can lower pH — a pattern similar to that observed in low-intensity fires and deeper soil horizons. Future studies could explore the full gradient of temperature–pH responses to better constrain local fire heating intensities, and investigate the specific chemistry driving these effects.

### Depth and heat flux interact to determine losses of preexisting PyC in subsequent fires

The self-evident observation that PyOM remains combustible is sometimes overlooked in discussions of its potential role as a C sink in terrestrial ecosystems. However, past studies have clearly reflected this: Doerr et al. (2018)^77^ found that higher fire intensity caused higher mass and PyC losses, and Tinkham et al. (2016)^78^ found that less PyC remained after each reburn. Thus, our findings of PyC losses during the reburn, and more C loss under high heat fluxes do not themselves come as a surprise. Our results are consistent with our hypotheses and past work – for all treatments, we observed C losses, and more C loss occurred under high heat fluxes when the PyOM was at the same depth (Fig. 6).

Perhaps the most important observations from this study are related to the interactions between depth and heat flux. Previous related studies have not studied depth as a variable, although Tinkham et al. (2016)^78^ suggested that burial depth > 3 cm in the mineral soil could prevent thermal degradation. However, in our experiment, at the depth of 5 cm, 40.1±1.8% and 2.6±0.6% of total C was lost under High HF and Low HF treatments, respectively (Table S7.2), indicating the interactive effects between exposure depth and heat treatment on protection from thermal degradation.

Regardless, though, C losses should be consistently expected to decrease with depth, with greatest – and in some cases, near-complete – losses at and near the surface (Fig. 6; Table S7.2). These findings underscore the critical importance of understanding interactions between changes in fire severity and fire return intervals, and burial rates of PyOM. If PyOM burial / translocation rates are sufficiently high so as to outpace fire return intervals – for example, moving down from O horizon to mineral soil within one year postfire – PyOM is likely to remain protected from losses in subsequent fires^42^. However, if fire regimes change such that fire frequency increases, or fire severity increases, natural burial rates may become insufficient to preserve preexisting PyOM, resulting in C losses^77,79^. Similarly, if the mechanisms driving burial rates change (e.g., due to changing precipitation, wind, or sediment deposition rates)^80^, critical losses in PyOM stocks may occur with future fire regimes. Conversely, if changes in burial rates or fire regimes interact to increase the critical differences between burial rates and fire return intervals or fire severity, then PyOM stocks have the potential to increase^4^. Modelling approaches may be required to help us resolve the range of scenarios expected for a given system or landscape^52^.

### Subsequent fires may further increase PyC losses through increased mineralizability and DOC production

Contrary to our hypothesis, we found that subsequent fires increased mineralizability (Fig. 7b) and overall decomposition rate (Fig. 8b) – so much so, that even the total C respired was greater than in the controls (Fig. 7a), despite the reduction in total C remaining in the burned treatments. These results underscore that, at certain burial depths and heat treatments, not only are there direct losses of C due to combustion, but the remaining C is even more susceptible to decomposition, further exacerbating potential net C losses from subsequent fires. Furthermore, these readily mineralizable fractions may induce positive priming (stimulation of the decomposition of other soil organic matter) after the addition of PyOM, further diminishing the effects of PyC on C sequestration over time^39,63^. Mineralized C was positively correlated with, and of a similar magnitude to, total DOC (Fig. 10a). This positive correlation is consistent with past findings that DOC is likely an important contributor to the readily mineralized fraction of the total C that is more readily decomposed by microbes^60^.

Although DOC represents a relatively small fraction of total PyC, it is important due to its mineralizability and also its potential translocation^81^. The release of dissolved PyC occurs slowly over time even when fire is absent^82^, but dissolved PyC is still thought to account for most of the vertical movement and burial of PyC in mineral soil^8,35^. Previous studies have observed that oxidation from fire increases the water solubility of PyC^8^, and other studies have observed more DOC leached from surface soil with lower-temperature heating (300 °C) than with higher temperatures (450 °C)^81^, which is consistent with our results. Deep dissolved PyC could form an important part of OM stocks in subsoil horizons, where half of total SOC stocks are located^83^, as dissolved PyC can be preferentially transported over other forms of DOC^17^. However, based on our results, without any immediate translocation mechanisms (such as rainfall), the majority of dissolved PyC may instead be mineralized by microbes rapidly after fire.

### Experimental limitations

Our experiments did not directly measure the properties of PyOM alone, but, rather, quantified them in the ground PyOM-sand samples. As discussed in the methods, we expect this would generally change the magnitude of some parameters, but not likely the relationships between the treatments. For example, our analyses of burned PyOM used ground samples, which means we neglected any effects of the particle size of PyOM, which would be expected to affect absolute respiration rates. We believe our sampling method did effectively recapture the full PyOM samples from the sand, given that the C loss fraction closely mirrored the mass loss fraction^53^ (Fig. S7.1; Table S7.1; Table S7.2) and that the mass of C in the retrieved control PyOM-sand samples is what we would predict from %C and mass of PyOM alone (Text S7.1).

Based on our uninoculated control incubation, we were unable to conclusively determine how much emitted CO_2_ is from inorganic C (Fig. S8.1, Table S8.1). There were leaks in the incubation jars, resulting in the saturation of KOH traps in nearly 1/3 of the samples, and we did not have sufficient PyOM materials to redo the incubation. Despite this, with the remaining measurements, we observe that some of the emitted CO_2_ is likely derived from inorganic C (inferred from observed CO_2_ emissions from uninoculated samples), but these emissions are consistently less than in the inoculated controls (Fig. 7a and Fig. S8.1). We also recognize that although we do not expect meaningful populations of microbes to exist in the ashed sand and PyOM, the PyOM-sand samples were not actively sterilized, so small populations of existing microbes could have been activated upon the addition of the sterile nutrient solution, also contributing to the CO_2_ emissions in the uninoculated control.

### Implications for land management and future research

Our results indicate that PyOM combustion is more likely under higher fire intensities and when PyOM remains at the soil surface. Fire management decisions can directly influence the fate of PyOM by affecting both likelihood of combustion and the opportunity for burial between fires.

In general, low-severity fires cause less disturbance to soils^9^, but may also be associated with higher frequency, and, hence, shorter time for PyOM burial between fires. Increased fire frequency could also potentially increase combustion of PyC and negatively influence long-term accumulation^8^. Management practices that increase tree and fuel density may elevate the risk of high-severity fires, especially under climate change^84,85^, thereby increasing the likelihood of PyC losses through deep combustion. Modeling approaches could help predict the net effect of changing fire regimes on soil C and PyC stocks, representing tradeoffs between PyC production and loss rates, as well as the dynamics of non-pyrogenic C.

If the goal is to increase or maintain soil PyC stocks, fire management should consider the factors that 1) affect heat flux to soils, including fuel loads^86^, fuel removal strategies, and potential effects of fires escaping; 2) govern PyOM burial, including typical fire return intervals and whether they will be sufficiently long for PyOM to move deeper into the soil. However, rates and mechanisms of PyOM incorporation are still poorly understood, highlighting a key need for future research across ecosystems and soil types. While carbon retention may be one goal of land management, it often intersects with others such as forest regeneration, wildfire risk mitigation, or habitat restoration. Clear communication among scientists, land managers, and policymakers is essential to align objectives and achieve co-benefits across fire-affected landscapes.

## Conclusions

Based on the results of our full-factorial experiment with different burial depths of PyOM and different heat flux profiles, we conclude that subsequent fires consume residual PyOM while also making the remaining PyOM more susceptible to microbial decomposition. We found that high heat flux and/or surface fires (HF_High_+Surface, HF_High_+1cm, and HF_Low_+Surface) resulted in large direct C losses through combustion. Intermediate heat exposure (HF_High_+5cm and HF_Low_+1cm) resulted in combustion losses along with increased DOC and mineralizability, which could lead to more complicated long-term effects: increases in the dissolved fraction of PyOM may make it more easily move downward into mineral soil and contribute to the persistent C pool in deep soil, but may also make it more readily decomposed by microbes. For the lowest heat flux and deepest burial (HF_Low_+5cm), most PyC was retained, and changes to C mineralization and DOC were minimal. Finally, pH, an important chemical property of PyOM, was subject to decreases at low-temperature heating but increases at higher temperatures. This work leaves many questions unanswered, including an expansion of this approach to encompass a broad range of fire regimes and ecosystems, to provide a holistic view of the impact of repeated fires on the global PyC cycle. Modeling may prove critical to predict the net effects of the complex interactions governing C persistence under repeated fires.

## Acknowledgements

This study was funded by the Department of Energy as a part of the project ‘Dissection of Carbon and Nitrogen Cycling in Post-Fire Soil Environments using a Genome-Informed Experimental Community’ DE-SC0020351. ML was also supported during part of this work by a UW-Madison Hatch grant. We would like to thank Roger Bohringer from Wilson State Forest Nursery for the donation of the jack pine tree. We also want to thank Troy Humphrey from Department of Soil and Environmental Sciences for helping cut and grind the tree branches to produce PyOM and Kelsey Kruger for additional support and help.

## Data Availability

Data and code associated with this project can be found at https://github.com/MengmengLuo/Fire-removes-preexisting-pyrogenic-organic-matter-from-theecosystem and the US Department of Energy ESS-DIVE database as Luo M; Yedinak K; Bourne K; Whitman T (2025): Heating effects on pyrogenic organic matter properties from a pyrocosm study in 2022. Dissection of Carbon and Nitrogen Cycling in Post-Fire Soil Environments using a Genome-Informed Experimental Community (DE-SC0020351). Dataset. doi:10.15485/3005739.

