## Supplemental Information for "Fire removes preexisting pyrogenic organic matter from the ecosystem through the mechanisms of both direct combustion and increasing mineralizability"

### 1. pH and segmented model

| Table S1.1. Segmented model summary | | | | |
| --- | --- | --- | --- | --- |
| Estimated Break-Point: x = 273 | | | | |
|  | Estimate | Std. Error | t-value | p-value |
| Intercept $\beta_{0}$ | 6.85 | 0.20 | 34.50 | <2 × 10^-16^ |
| $\beta_{1}$ | -0.0014 | 0.0011 | -1.31 | 0.20 |
| $U_{1}$ | 0.0040 | 0.0011 | 3.56 | NA |
| Residual standard error: 0.18 on 26 degrees of freedom  Multiple R-Squared: 0.78, Adjusted R-squared: 0.75 | | | | |

#### Text S1.1. Segmented model equation and description

The model equation for pH is:

$$pH{=\beta}_{0}-\beta_{1}\cdot T_{Peak}+U_{1}\cdot[T_{Peak}-273]$$

The breakpoint of the segmented model is 273. The notation $\left[ T_{Peak}-273 \right]$ means that if $T_{Peak}$<273, then $\left[ T_{Peak}-273 \right]$ =0, which means this term does not apply to the model. If $T_{Peak}$>273, then $\left[ T_{Peak}-273 \right]=(T_{Peak}-273)$, which means this term applies to the model.

From this equation, $\beta_{0}$, the intercept for $T_{Peak}$<273, is 6.85, based on the p value, the intercept is significantly different from zero. $\beta_{1}$ is the slope for $T_{Peak}$<273, which is -0.0014. Based on the p value, the slope is not significantly different from zero, which means that for $T_{Peak}$<273, peak temperature does not significantly affect the pH for PyOM. $U_{1}$ is the change in slope after the break point, which is 0.0040. So following is the functions for this segmented model (also indicated in Fig. 4b):

$$pH=6.85-0.0014\cdot T_{Peak} (T_{Peak}<273)$$

$$pH=5.75+0.0026\cdot T_{Peak} (T_{Peak}>273)$$

Because the p value was not displayed for the slope after the breakpoint, we checked the 95% confidence level (CI) to see if the CI includes zero. Because the 95% CI is (199.10, 347.05), which means when $T_{Peak}$>273, peak temperature has a significant positive effect on pH.

| Table S1.2. ANOVA statistics for pH | | | | | |
| --- | --- | --- | --- | --- | --- |
| Sources | df | Sum of Squares | Mean Squares | F statistic | p-value |
| Heat Flux | 2 | 0.86 | 0.43 | 51.86 | < 2.51 × 10^-11^ |
| Depth | 2 | 0.73 | 0.37 | 43.79 | < 2.28 × 10^-10^ |
| Heat Flux × Depth | 4 | 2.08 | 0.52 | 62.45 | < 1.07 × 10^-15^ |
| Residuals | 36 | 0.30 | 0.0083 |  |  |

### 2. Remaining C in PyOM-sand samples

| Table S2.1. ANOVA statistics for total C (mg per sample) | | | | | |
| --- | --- | --- | --- | --- | --- |
| Sources | df | Sum of Squares | Mean Squares | F statistic | p-value |
| Heat Flux | 2 | 2953713 | 1476856 | 8580 | < 2 × 10^-16^ |
| Depth | 2 | 1269206 | 634603 | 3687 | < 2 × 10^-16^ |
| Heat Flux × Depth | 4 | 717632 | 179408 | 1042 | < 2 × 10^-16^ |
| Residuals | 36 | 6197 | 172 |  |  |

### 3. Mineralized C

| Table S3.1. ANOVA statistics for CO_2_-C (mg per sample) | | | | | |
| --- | --- | --- | --- | --- | --- |
| Sources | df | Sum of Squares | Mean Squares | F statistic | p-value |
| Heat Flux | 2 | 7.87 | 3.94 | 7.69 | 0.0017 |
| Depth | 2 | 71.52 | 35.76 | 69.89 | 5.99 × 10^-13^ |
| Heat Flux × Depth | 4 | 154.93 | 38.73 | 75.70 | < 2 × 10^-16^ |
| Residuals | 35 | 17.91 | 0.51 |  |  |

### 4. C mineralizability

| Table S4.1. ANOVA statistics for C mineralizability (mg CO_2_-C × mg^-1^ C) | | | | | |
| --- | --- | --- | --- | --- | --- |
| Sources | df | Sum of Squares | Mean Squares | F statistic | p-value |
| Treatments | 6 | 0.0014 | 2.29 × 10^-4^ | 8.33 | 3.16 × 10^-5^ |
| Residuals | 28 | 0.00077 | 2.75 × 10^-5^ |  |  |

### 5. Readily degradable carbon fraction (coefficient a) and degradation rate (coefficient b)

| Table S5.1. Linear model output for mineralizability vs. readily degradable C fraction (coefficient a) | | | | |
| --- | --- | --- | --- | --- |
| Residuals: | | | | |
| Min | 1Q | Median | 3Q | Max |
| -0.0094 | -0.00095 | -0.00060 | 0.0020 | 0.0046 |
|  | Estimate | Std. Error | t-value | p-value |
| Intercept | 0.0015 | 0.00078 | 1.99 | 0.055 |
| Slope | 0.74 | 0.046 | 15.83 | < 2 × 10^-16^ |
| Residual standard error: 0.002595 on 32 degrees of freedom  Multiple R-squared: 0.8868, Adjusted R-squared: 0.8832  F-statistic: 250.6 on 1 and 32 DF, p-value: < 2.2e-16 | | | | |

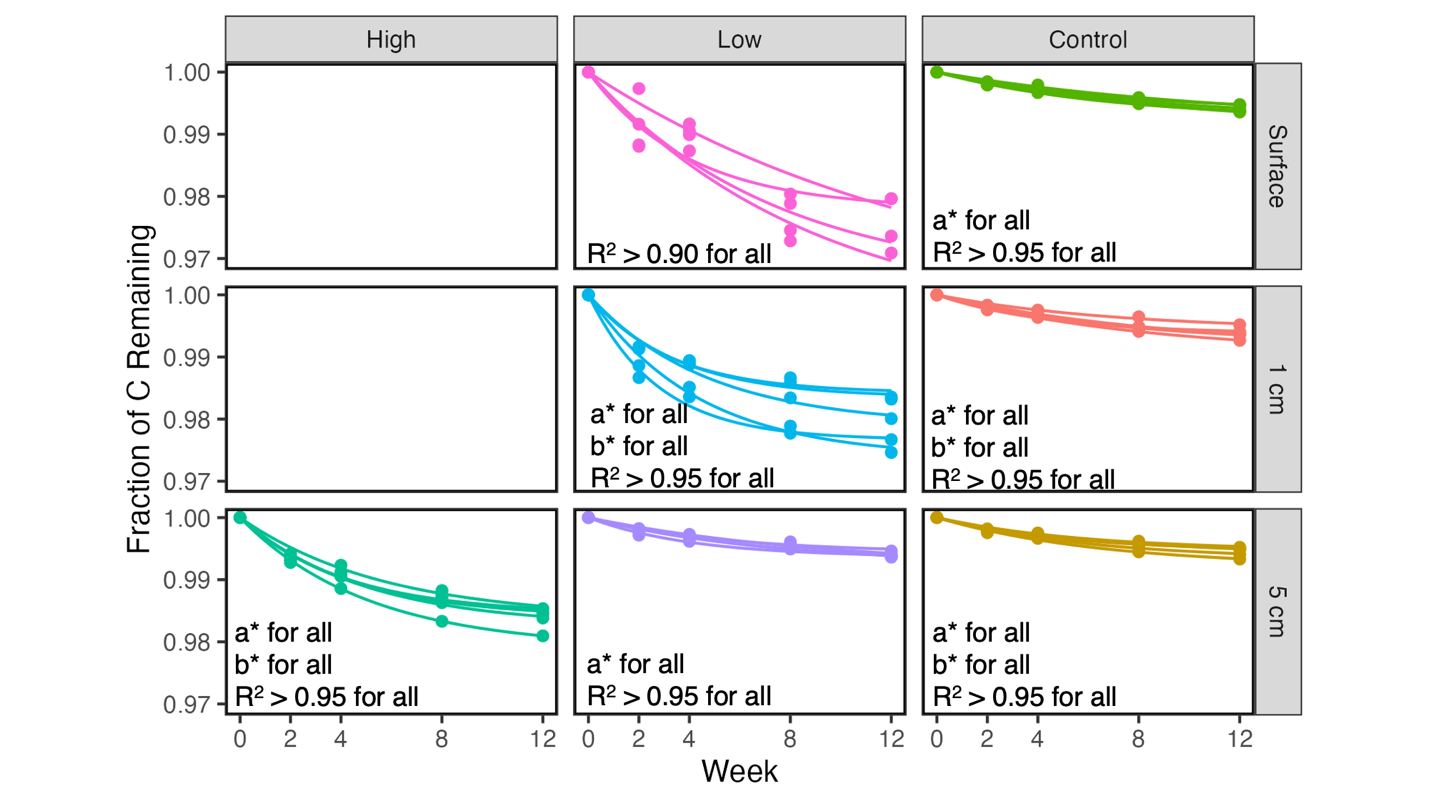

Figure S5.1. One-pool decay models fitting with fraction C remaining over 12 weeks’ incubation. n = 5, excluded HF_High_+Surface and HF_High_+1cm for unquantifiable total C, and one outlier in HF_Low_+Surface (iterative *z* = -3.93 for coefficient *a*; iterative *z* = 10.30 for coefficient *b*).

| Table S5.2. ANOVA statistics for readily decomposable C fraction (coefficient a) | | | | | |
| --- | --- | --- | --- | --- | --- |
| Sources | df | Sum of Squares | Mean Squares | F statistic | p-value |
| Treatments | 6 | 0.0028 | 0.00047 | 37.97 | 6.31 × 10^-12^ |
| Residuals | 27 | 0.00033 | 0.000012 |  |  |

| Table S5.3. ANOVA statistics for C decomposition rate (coefficient *b*, week^-1^) | | | | | |
| --- | --- | --- | --- | --- | --- |
| Sources | df | Sum of Squares | Mean Squares | F statistic | p-value |
| Treatments | 6 | 0.088 | 0.015 | 6.65 | 0.00021 |
| Residuals | 27 | 0.060 | 0.0022 |  |  |

#### Table S5.4. Statistics for one-pool decay model

| HF | Depth | a±SE | b±SE | P value for a | P value for b | R^2^ |
| --- | --- | --- | --- | --- | --- | --- |
| High | 5 cm | 0.017±0.00072 | 0.20±0.020 | 0.00016 | 0.0019 | 0.997 |
| High | 5 cm | 0.016±0.0010 | 0.23±0.036 | 0.00052 | 0.0077 | 0.990 |
| High | 5 cm | 0.016±0.0017 | 0.17±0.038 | 0.0023 | 0.020 | 0.986 |
| High | 5 cm | 0.021±9.71×10^-5^ | 0.20±0.0021 | 2.18×10^-7^ | 2.63×10^-6^ | 1.000 |
| High | 5 cm | 0.015±0.0015 | 0.24±0.058 | 0.0018 | 0.025 | 0.976 |
| Low | Surface | 0.034±0.013 | 0.13±0.095 | 0.078 | 0.25 | 0.910 |
| Low | Surface | 0.041±0.0099 | 0.11±0.047 | 0.026 | 0.094 | 0.977 |
| Low | Surface | 0.022±0.0042 | 0.25±0.13 | 0.013 | 0.14 | 0.899 |
| Low | Surface | 0.038±0.027 | 0.070±0.070 | 0.26 | 0.39 | 0.948 |
| Low | 1 cm | 0.021±0.0017 | 0.21±0.042 | 0.0012 | 0.014 | 0.986 |
| Low | 1 cm | 0.016±0.0012 | 0.31±0.072 | 0.0010 | 0.023 | 0.973 |
| Low | 1 cm | 0.026±0.0019 | 0.23±0.043 | 0.00089 | 0.013 | 0.986 |
| Low | 1 cm | 0.023±0.0011 | 0.36±0.055 | 0.00021 | 0.0062 | 0.989 |
| Low | 1 cm | 0.017±0.0011 | 0.29±0.055 | 0.00067 | 0.014 | 0.982 |
| Low | 5 cm | 0.0057±0.00081 | 0.18±0.058 | 0.0059 | 0.050 | 0.970 |
| Low | 5 cm | 0.0064±0.00050 | 0.24±0.049 | 0.0010 | 0.016 | 0.983 |
| Low | 5 cm | 0.0074±0.0014 | 0.14±0.052 | 0.014 | 0.072 | 0.971 |
| Low | 5 cm | 0.0075±0.00036 | 0.14±0.013 | 0.00023 | 0.0015 | 0.998 |
| Low | 5 cm | 0.0080±0.00091 | 0.11±0.020 | 0.0031 | 0.013 | 0.995 |
| Control | Surface | 0.0082±0.0019 | 0.086±0.030 | 0.023 | 0.065 | 0.989 |
| Control | Surface | 0.0076±0.00033 | 0.15±0.012 | 0.00019 | 0.0013 | 0.998 |
| Control | Surface | 0.0092±0.0018 | 0.098±0.030 | 0.013 | 0.046 | 0.989 |
| Control | Surface | 0.0064±0.00099 | 0.14±0.042 | 0.0075 | 0.043 | 0.981 |
| Control | Surface | 0.0075±0.00074 | 0.12±0.021 | 0.0020 | 0.010 | 0.995 |
| Control | 1 cm | 0.0085±0.00034 | 0.11±0.0080 | 0.00015 | 0.00073 | 0.999 |
| Control | 1 cm | 0.0095±0.00075 | 0.12±0.017 | 0.0011 | 0.0058 | 0.997 |
| Control | 1 cm | 0.0079±0.00023 | 0.14±0.0076 | 5.12×10^-5^ | 0.00035 | 0.999 |
| Control | 1 cm | 0.0057±0.00081 | 0.14±0.037 | 0.0059 | 0.034 | 0.985 |
| Control | 1 cm | 0.0063±0.00017 | 0.23±0.015 | 4.19×10^-5^ | 0.00062 | 0.998 |
| Control | 5 cm | 0.0053±0.00059 | 0.17±0.040 | 0.0029 | 0.024 | 0.985 |
| Control | 5 cm | 0.0060±0.00053 | 0.15±0.027 | 0.0015 | 0.011 | 0.993 |
| Control | 5 cm | 0.0079±0.00070 | 0.15±0.027 | 0.0015 | 0.011 | 0.993 |
| Control | 5 cm | 0.0068±0.00043 | 0.16±0.022 | 0.00056 | 0.0047 | 0.996 |
| Control | 5 cm | 0.0053±0.00066 | 0.21±0.060 | 0.0040 | 0.042 | 0.970 |

### 6. Dissolved organic and inorganic carbon

| Table S6.1. Linear model output for CO_2_-C vs. DOC | | | | |
| --- | --- | --- | --- | --- |
| Residuals: | | | | |
| Min | 1Q | Median | 3Q | Max |
| -2.18 | -0.61 | -0.11 | 0.50 | 2.84 |
|  | Estimate | Std. Error | t-value | p-value |
| Intercept | -2.21 | 0.46 | -4.77 | 2.27 × 10^-5^ |
| Slope | 1.35 | 0.093 | 14.59 | < 2 × 10^-16^ |
| Residual standard error: 0.99 on 42 degrees of freedom  Multiple R-squared: 0.84, Adjusted R-squared: 0.83  F-statistic: 212.9 on 1 and 42 DF, p-value: < 2.2e-16 | | | | |

| Table S6.2. Linear model output for pH vs. DOC | | | | |
| --- | --- | --- | --- | --- |
| Residuals: | | | | |
| Min | 1Q | Median | 3Q | Max |
| -0.36 | -0.080 | 0.033 | 0.073 | 0.38 |
|  | Estimate | Std. Error | t-value | p-value |
| Intercept | 7.56 | 0.066 | 114.64 | < 2 × 10^-16^ |
| Slope | -0.16 | 0.013 | -12.24 | 1.35 × 10^-15^ |
| Residual standard error: 0.14 on 43 degrees of freedom  Multiple R-squared: 0.78, Adjusted R-squared: 0.77  F-statistic: 149.7 on 1 and 43 DF, p-value: 1.35e-15 | | | | |

| Table S6.3. Linear model output for pH vs. DIC | | | | |
| --- | --- | --- | --- | --- |
| Residuals: | | | | |
| Min | 1Q | Median | 3Q | Max |
| -0.54 | -0.12 | 0.029 | 0.095 | 0.62 |
|  | Estimate | Std. Error | t-value | p-value |
| Intercept | 6.12 | 0.10 | 60.50 | < 2 × 10^-16^ |
| Slope | 1.29 | 0.18 | 7.072 | 1.01 × 10^-8^ |
| Residual standard error: 0.21 on 43 degrees of freedom  Multiple R-squared: 0.54, Adjusted R-squared: 0.53  F-statistic: 50.01 on 1 and 43 DF, p-value: 1.01e-08 | | | | |

| Table S6.4. ANOVA statistics for DOC (mg per sample) | | | | | |
| --- | --- | --- | --- | --- | --- |
| Sources | df | Sum of Squares | Mean Squares | F statistic | p-value |
| Heat Flux | 2 | 20.10 | 10.05 | 65.06 | 1.11 × 10^-12^ |
| Depth | 2 | 32.48 | 16.24 | 105.12 | 9.31 × 10^-16^ |
| Heat Flux × Depth | 4 | 58.87 | 14.72 | 95.27 | < 2 × 10^-16^ |
| Residuals | 36 | 5.56 | 0.16 |  |  |

| Table S6.5. ANOVA statistics for DIC (mg per sample) | | | | | |
| --- | --- | --- | --- | --- | --- |
| Sources | df | Sum of Squares | Mean Squares | F statistic | p-value |
| Heat Flux | 2 | 0.29 | 0.15 | 4.65 | 0.016 |
| Depth | 2 | 0.52 | 0.26 | 81.92 | 3.99 × 10^-14^ |
| Heat Flux × Depth | 4 | 0.63 | 0.16 | 50.06 | 3.19 × 10^-14^ |
| Residuals | 36 | 0.11 | 0.0032 |  |  |

### 7. Estimated carbon loss fraction and percentage carbon in the original PyOM

#### Text S7.1. C Loss Fraction description

C Loss Fraction ($C_{L}$) was estimated using Equation S1,

$$\begin{aligned} C_{L} = \frac{C_{U}\times M_{U}- C_{B}\times M_{B}}{C_{U}\times M_{U}} \#\left( S1 \right) \end{aligned}$$

where C_B_ is the mass percent C of the mass of each burned sample, and M_B_ is the sample mass of each burned sample. C_U_ is the mean value of the mass percent C for the mean mass of the unburned samples (n = 5) at the same corresponding depth. The linear model fit indicates that the mass loss fraction of PyOM (reported in Luo et al., 2025) closely mirrors the fractional C loss (Fig. 4b, R^2^=0.98).

For the 15 controls, we calculated theoretically how much C% should be in each original PyOM sample, using equation S2, where *C_L_* is carbon loss fraction as calculated in Equation S1, $M$ is the final sample mass (~80g), and $m_{PyOM}$ is the mass of PyOM in each sample (~1g).

$$\begin{aligned} C_{E} =\frac{(1- C_{L})\cdot M}{m_{PyOM}}\#\left( S2 \right) \end{aligned}$$

We used the Wilcox test to test if the data for the 15 controls (n = 15) is significantly different from the actual C% in the PyOM as measured directly (n = 3). The estimated C concentration for the controls was not significantly different from the direct measurement for C in original PyOM (p = 0.65), which indicates that our methods of extraction after burial and subsequent processing likely capture all or most of the PyOM-C.

| Table S7.1. Linear model output for C loss fraction vs. mass loss fraction | | | | |
| --- | --- | --- | --- | --- |
| Residuals: | | | | |
| Min | 1Q | Median | 3Q | Max |
| -0.085 | -0.023 | -0.0060 | 0.026 | 0.10 |
|  | Estimate | Std. Error | t-value | p-value |
| Intercept | 0.040 | 0.019 | 2.12 | 0.043 |
| Slope | 0.97 | 0.026 | 36.97 | < 2 × 10^-16^ |
| Residual standard error: 0.052 on 28 degrees of freedom  Multiple R-squared: 0.98, Adjusted R-squared: 0.98  F-statistic: 1367 on 1 and 28 DF, p-value: < 2.2e-16 | | | | |

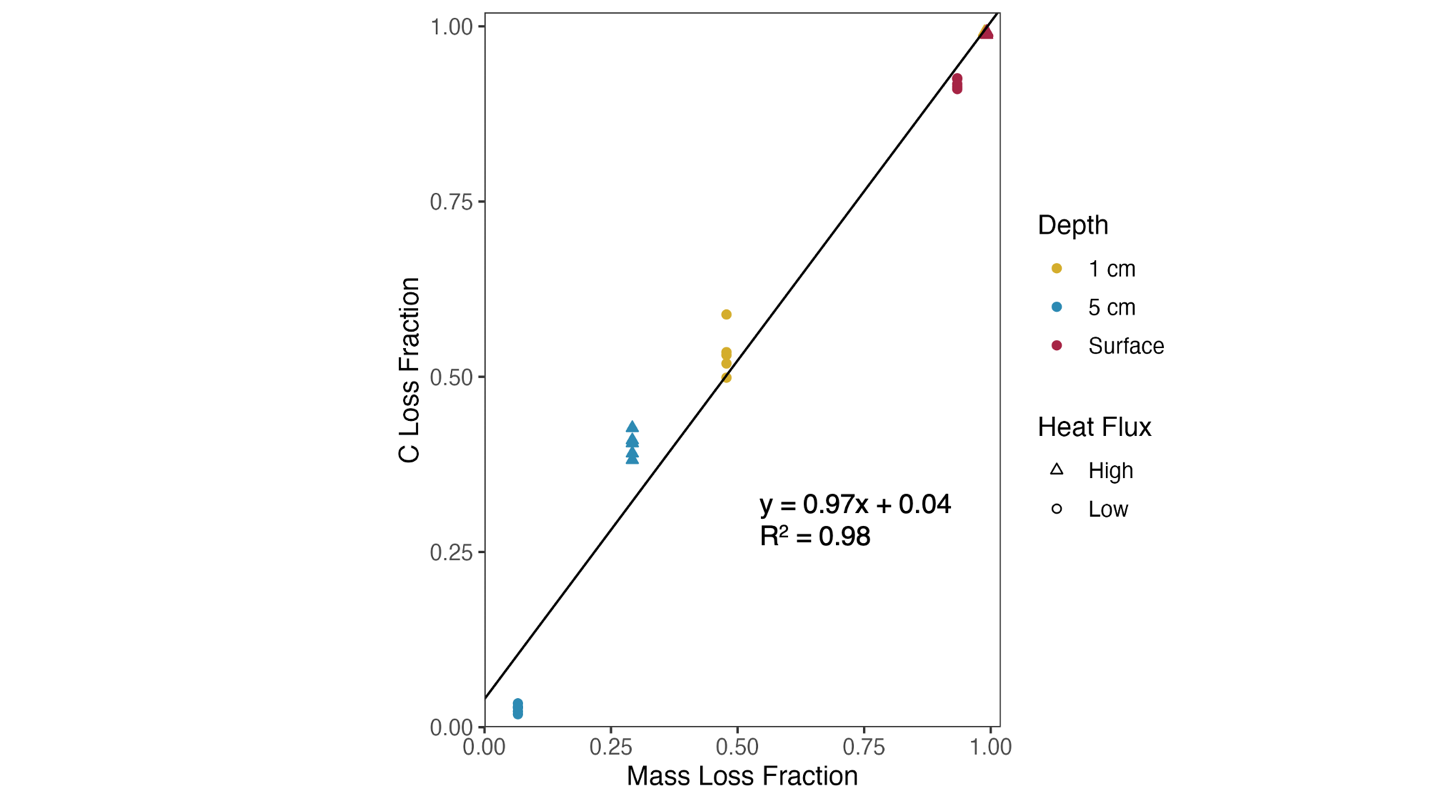

Figure S7.1. C loss fraction vs. PyOM mass loss fraction (line indicates linear model fit; p < 0.05 for both slope and intercept; R^2^=0.98)

| Table S7.2. C loss fraction for all treatments (n = 5; controls not included); Different superscript letters (ANOVA, Tukey’s HSD) indicate significant differences across all treatments | | | |
| --- | --- | --- | --- |
| Heat Flux | Depth | C Loss Fraction (%) | SD (%) |
| High | Surface | 99.05^a^ | 0.16 |
| High | 1cm | 99.05^a^ | 0.13 |
| High | 5cm | 40.09^d^ | 1.76 |
| Low | Surface | 91.78^b^ | 0.69 |
| Low | 1cm | 53.22^c^ | 3.37 |
| Low | 5cm | 2.57^e^ | 0.60 |

### 8. Blank incubation details and limitations

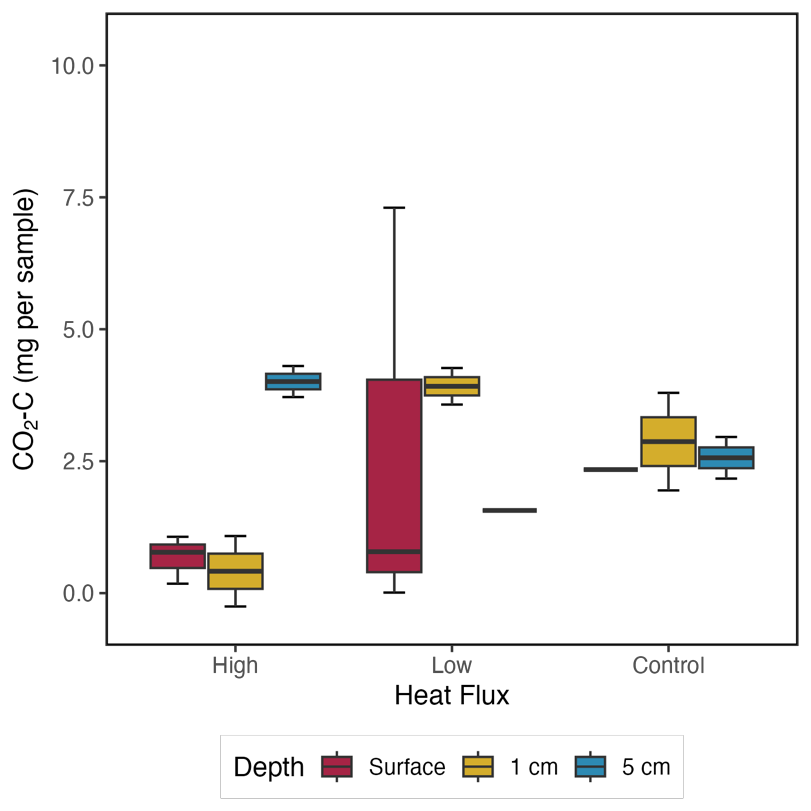

Figure S8.1. C loss as CO_2_ during 12-week incubation with no inoculants (n = 1 for Control+Surface and HF_Low_+5cm; n = 2 for HF_High_+1cm, HF_High_+5cm, HF_Low_+1cm, , Control+1cm, Control+5cm; n = 3 for HF_High_+Surface and HF_Low_+Surface)

| Table S8.1. ANOVA statistics for CO_2_-C (blank incubation, mg per sample) | | | | | |
| --- | --- | --- | --- | --- | --- |
| Sources | df | Sum of Squares | Mean Squares | F statistic | p-value |
| Heat Flux | 2 | 6.78 | 3.39 | 0.85 | 0.46 |
| Depth | 2 | 4.46 | 2.23 | 0.56 | 0.59 |
| Heat Flux × Depth | 4 | 16.69 | 4.17 | 1.05 | 0.44 |
| Residuals | 9 | 35.82 | 3.98 |  |  |
